# The Wsp intermembrane complex mediates metabolic control of the swim-attach decision of *Pseudomonas putida*

**DOI:** 10.1101/2020.02.05.934950

**Authors:** Ángeles Hueso-Gil, Belén Calles, Víctor de Lorenzo

## Abstract

*Pseudomonas putida* KT2440, a microorganism of interest for biotechnological purposes, is one amongst the many bacteria that attach to surfaces and produce biofilm. Although other mechanisms that contribute to this decision have been studied until now, a 7-genes-operon with a disposition and homology shared with the *wsp* operon in *Pseudomonas aeruginosa* remained to be investigated. In this work, we characterized the function of *P. putida wsp* operon by the combination of deletion mutants with complementations with *P. aeruginosa*’s genes and with deletions of 3 other genes: the genes that code for the transcription factors *fleQ* and *fleN* and the flagellar movement regulator, *fglZ*. Examining mutant behaviour at 6 and 24 hours under three different carbon regimes (citrate, glucose and fructose) we saw that this complex carries out a similar function in both *Pseudomonas*. In *P. putida*, the key components are WspR, a protein that harbours the domain for producing c-di-GMP, and WspF, which controls its activity. Transformation with the equivalent proteins of *P. aeruginosa* had a significant impact on of *P. putida* mutant phenotypes and could complement their functions under some conditions. These results contribute to the deeper understanding of the complex element network that regulate lifestyle decision in *P. putida*

## INTRODUCTION

The environmental bacterium *Pseudomonas putida* KT2440 has is growingly receiving attention a a suitable chassis for a suite or biotechnological and environmental applications. In this context, the characterization of the mechanisms that trigger the swim or attach decision are decisive for e.g. developing catalytic biofilms. Its close relative *Pseudomonas aeruginosa* PAO1 has been a biofilm model system in Gram-negative bacteria because of the pathogenic consequences of its attachment to prostheses and its effects in lung colonization. The intermembrane Wsp complex present in *P. aeruginosa* and other pseudomonads modulates attachment controlling the formation of c-di-GMP, the main secondary messenger that regulates the attachment-swimming decision (Simm *et al.*, 2004; Hickman *et al.*, 2005). However, the starting signal that activates the complex is not very well defined yet, although it is probably directly related to physical contact. Amongst the seven protein Wsp intermembrane complex from *P. aeruginosa* (Figure 1), structural component WspA is usually located in dynamic clusters in the laterals of bacterial cell surface and, under certain circumstances such as growth on agar surface, it transduces a signal through the entire complex with the special intervention of WspE (Guvener and Harwood, 2007; O’Connor *et al.*, 2012). WspR, another protein in the complex, contains the enzymatic domain GGDEF, able to form c-di-GMP (Simm *et al*., 2004). Through WspF regulation, WspR gets phosphorylated by WspE and shifts its equilibrium to the formation of c-di-GMP. In the absence of WspF, WspR from *P. aeruginosa* shows high cyclase activity, increasing c-di-GMP amounts in the cell and leading, consequently, to superior biofilm formation. Therefore, WspF is a regulator by repression of WspR activity (Guvener and Harwood, 2007; Huangyutitham *et al.*, 2013). According to current models, WspR is a cytoplasmic low-activity monomer that changes its conformation upon phophorilation, forming highly-active tetramers (Huangyutitham *et al*., 2013). Apart from this regulation, WspR activity is repressed by c-di-GMP molecule, which binds to an allosteric site in the GGDEF domain and provokes the dissolution of the clusters and a relaxation of the tetramers leading them to an inhibited WspR dimer form (De *et al.*, 2008; Huangyutitham *et al*., 2013).

**Figure 1:**
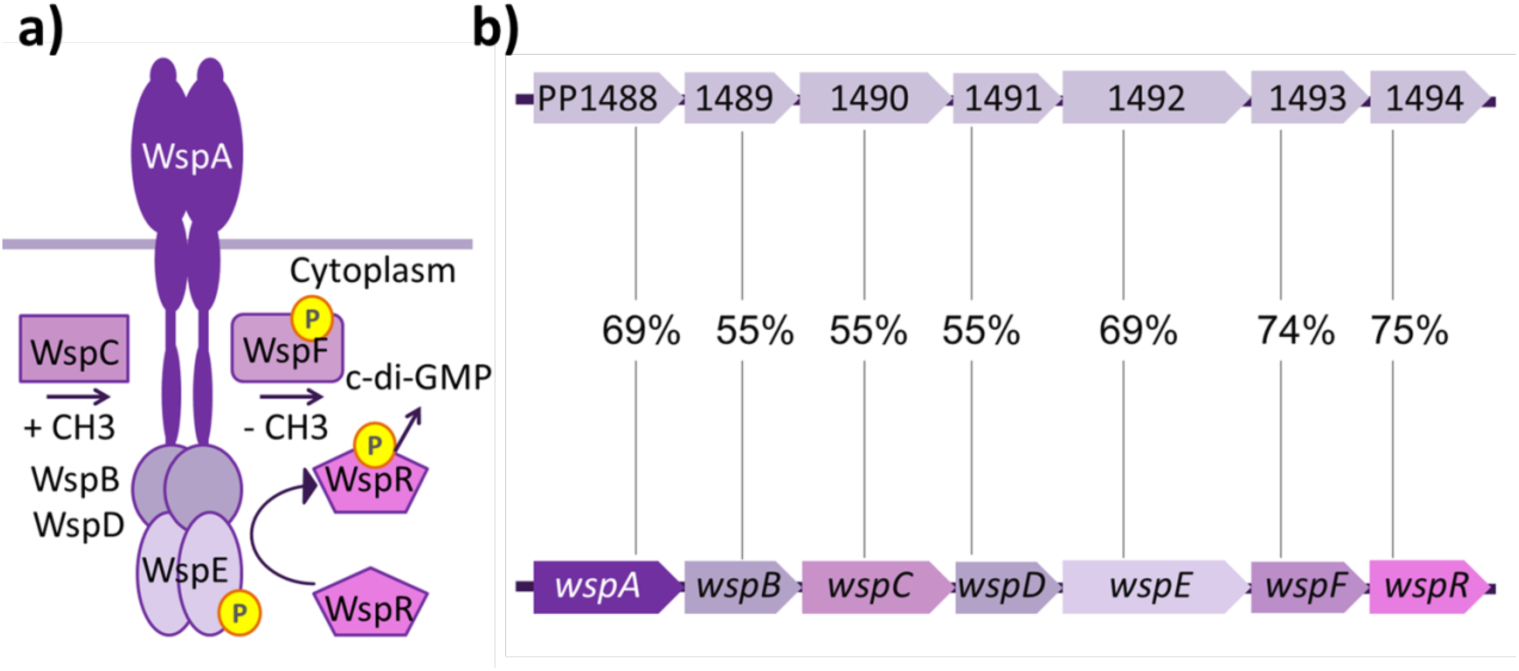
Wsp complex in *Pseudomonas* sp. **(a)** Model proposed for the structure of Wsp protein complex from *P. aeruginosa* (Guvener and Harwood, 2007). *P. putida*’s complex probably shows the same disposition. **(b)** Comparison between the seven genes of Wsp operon form *P. aeruginosa* (lower panel) and the putative syntenic operon found in *P putida* (upper panel). Identity percentage per gene is shown.

Another c-di-GMP-responsive transcription factor (TF) involved in biofilm formation and motility is the master regulator FleQ, that is also related to other functions, including iron homeostasis (Blanco-Romero *et al.*, 2018) and the control of the type 6 secretion system. (T6SS; (Wang *et al.*, 2018)). In *P. putida* in conditions of low c-di-GMP, FleQ can interact with σ^54^ to regulate a high number of promoters related to motility and flagella. However, when c-di-GMP concentration is high, this master regulator forms dimers and interacts with σ^70^ to activate or repress transcription of biofilm or flagella-related genes, respectively. In general, c-di-GMP binding to FleQ usually antagonizes its repressor activity, transforming this factor into an activator for the same promoter. This TF is so important in lifestyle decisions that a *P. putida* strain with *fleQ* deleted resulted in both non-motile and biofilm impaired bacteria (Jimenez-Fernandez *et al.*, 2016; Molina-Henares *et al.*, 2017). FleQ can also interact with another TF, FleN. In *P. aeruginosa*, FleN acts as an auxiliary element and antagonizes the FleQ dual function activator-repressor for the regulation of certain genes and operons (Dasgupta *et al.*, 2000; Baraquet *et al.*, 2012). In *P. putida*, FleN and FleQ also usually play opposite roles, with some exceptions such as their synergistic role in regulating transcription of the *bcs* biofilm operon and *lapA* adhesin (Nie *et al.*, 2017).

An additional actor of biofilm formation in pseudomonads is FlgZ (also called YcgR in *E. coli*). This is not a TF but a flagellar function regulator containing a PilZ domain, which can bind c-di-GMP (Amikam and Galperin, 2006). *flgZ* transcription depends on FleQ and FliA. In *E.coli*, YcgR regulates swimming velocity based on c-di-GMP levels of the cell (Ryjenkov *et al.*, 2006; Boehm *et al.*, 2010; Martinez-Granero *et al.*, 2014; Baker *et al.*, 2016). In the presence of c-di-GMP, it tends to dimerize and interacts with Mot components of the flagellar stator in addition to the motor and regulates its movement, but, at least in *Pseudomonas fluorescens*, FlgZ does this independent of c-di-GMP under the conditions tested. However, it does participate in the formation of biofilm mediated by c-di-GMP produced by WspR and degraded by the esterase BifA. In *P. putida* its presence is associated to a reduction of flagellar movement (Martinez-Granero *et al*., 2014).

On the basis of the above, we studied the *wsp* orthologous operon in *P. putida* by making individual and block-deletion mutants, in order to determine the importance of the similarities between this complex and *P. aeruginosa*’s and to decide how we could take advantage of its function in order to control biofilm formation. We also studied several mutant phenotypes in three different sole carbon sources, as the nature of them influences extracellular polysaccharide (EPS) building blocks used for biofilm formation (Reeves *et al.*, 1996; Ramos *et al.*, 2001; Wan *et al.*, 2018). The results below not only expose the similarities with the P. aeruginosa’s Wsp system of reference, but it also reveals a striking dependence of biofilm formation on growth conditions and metabolic status.

## RESULTS AND DISCUSSION

### Modulation of Pseudomonas putida attachment to surfaces through the Wsp complex

First of all, we searched for a region in *P. putida*’s genome that showed homology with the operon of *P. aeruginosa*’s *wsp*. Using the MicroScope platform from GenoScope (Évry, France), we found a set of genes from PP_1488 to PP_1494 *loci* that showed a significant synteny with the corresponding genes in *P. aeruginosa*, indicating a potentially shared evolutionary origin (Figure 1b). To study the role of the operon in biofilm formation we made nine clean deletions from ATG to stop codon, as indicated in the Materials and Methods section: one of them comprised the whole operon (*wsp*) while seven of them were deletions per each ORF (*wspA, wspB, wspC, wspD, wspE, wspF, wspR*) in order to study their individual effect. Finally, one more double mutant was made for the duo *wspF wspR*, since this region encodes the enzymatic activity and its regulatory protein. With this mutant collection in hand we then checked the phenotype of the different strains in terms of biofilm formation, colony morphology and swimming ability. Focusing on biofilm formation assays of Figure 2a and c, we found some differences with respect to three variables: between mutants, along time and regarding carbon source. As we can see, the general overview showed that carbon regime was an important factor for biofilm build up. Wild type *P. putida* KT2440 (*wt*) and the rest of the mutants showed a different biofilm profile in media supplemented with 3 types of sugar as a sole carbon source. For example, in glucose or fructose was able to form more biofilm than in citrate. For other side, the deletion mutant for the whole *wsp* operon produced similar amount of biofilm than *wt* cultures, indicating that the Wsp complex is keeping the cyclase activity of WspR in a low state in the conditions tested.

**Figure 2:**
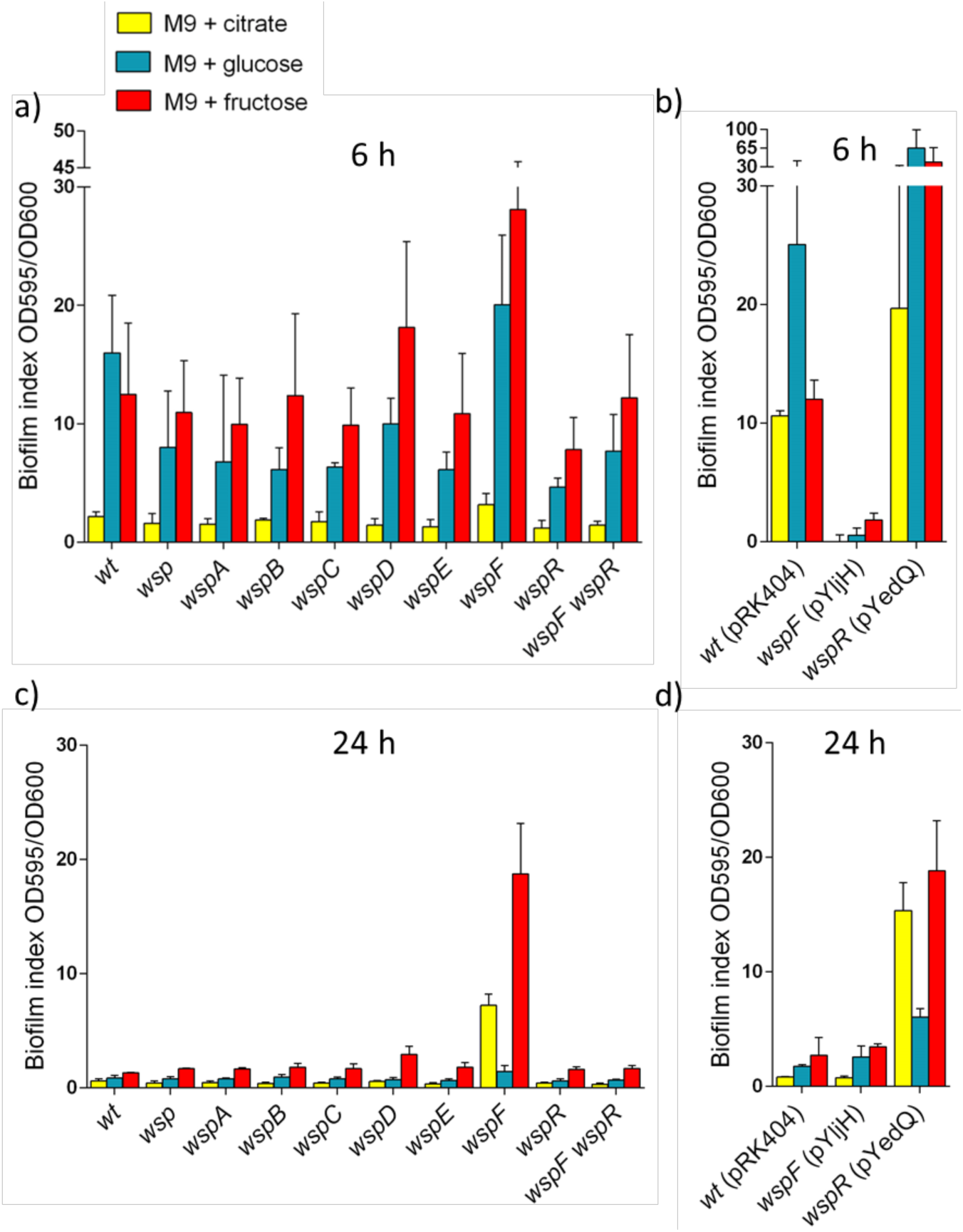
Biofilm assay for *P. putida wsp* mutants and their transformations with YedQ and YljH. All strains were cultured in M9 media supplemented with citrate (yellow), glucose (blue) or fructose (red) as sole carbon sources, respectively. **(a)** Biofilm was measured for *P. putida wt* and *wsp* mutants in the three carbon sources at 6 h of incubation as indicated. **(b)** Biofilm levels for the *wspF* strain transformed with the c-di-GMP cyclase YedQ carried by pYedQ plasmid and *wspR* transformed with the c-di-GMP esterase YljH carried by pYljH plasmid was checked for the three carbon sources after 6 h of incubation. The action of YljH is expected to reduce the biofilm formed by *wspF* mutant while the action of YedQ could increase the biofilm formation of the *wspR* deletion. **(c)** Biofilm levels for *P. putida wt* and *wsp* mutants at 24 h. **(d)** Biofilm measurement for *wspF* strain harbouring pYedQ plasmid and *wspR* bearing pYljH plasmid cultured for 24 h.

Considering time as another variable, we also noticed a general decrease in biofilm at 24 h, something that happened to the *wt* and to most of the mutants too. But, interestingly, although the *wspF* mutant formed slightly more biofilm than the rest of the deletions in all carbon sources tested at 6 h of incubation, those differences noticeably increased at 24 h due to distinct fluctuations between mutants. For example, in M9 citrate medium, the *wspF* mutant doubled biofilm formation from 6 h to 24 h, while the rest of the mutants decreased it slightly, contributing then to produce a higher difference between this deleted strain and the rest. Regarding M9 fructose medium, the *wspF* mutant kept a similar biofilm index from 6 h to 24 h, whereas the rest of the strains showed drops in production. In M9 glucose medium, this decline was specially intense affecting also to *wspF* mutant biofilm levels. Although biofilm fluctuations in time are inherent to *P. putida*, as *wt* levels showed, our results indicated that in the absence of WspF, WspR activity was not well regulated, leading to a longer in time c-di-GMP synthesis and biofilm production, but with some particularities depending on the carbon regime. On the contrary, *wspR* mutant was less able to form biofilm in some of the conditions tested (for example, at 6 h for glucose and fructose). This is consistent with the hypothesis that the activity of WspR, a protein with a GGDEF domain, is controlled by the phosphorylation regulated by WspF, as occurs in *P. aeruginosa* (Guvener and Harwood, 2007). According to this hypothesis for *P. putida*, in the absence of WspF, WspR would become phosphorylated at a higher rate by WspE such that it would continuously produce c-di-GMP and biofilm, while in the absence of WspR, the lower amount of c-di-GMP present in the bacteria would make it less able to form biofilm. For other side, the double mutant *wspF wspR* showed a similar biofilm production compared to the simple mutant *wspR*, as the deletion of the cyclase protein has a dominant phenotype over the deletion of its regulator.

### The interplay Wsp/ c-di-GMP / biofilm formation

To check whether the differences in biofilm formation between *wspF* and *wspR* mutants describe above were caused by c-di-GMP levels and not through other effectors, we transformed the *wspF* strain with plasmid pYljH, which encoded the YljH diguanylate esterase of *Escherichia coli*. We also transformed the *wspR* mutant with plasmid pYedQ carrying the YedQ diguanylate cyclase also from *E. coli*. Both proteins were expressed constitutively. We measured and compared the results between these new *wspF* and *wspR* strains bearing pYljH and pYedQ respectively, and *wt* strain with the empty plasmid, backbone pRK404, and we found that YljH esterase could notably decrease biofilm levels of *wspF* mutant at 6 h and 24 h (Figure 2b and d). In contrast, YedQ cyclase could significantly increase biofilm levels in the *wspR* mutant. Taking a closer look at the three carbon sources, the *wspR* strain transformed with YedQ enzyme could emulate the *wspF* biofilm phenotype in fructose at 6 and 24 h, while in citrate and glucose it would exceed it. However, the *wspF* strain with YljH reduced biofilm levels below those of the *wspR* mutant at 6 h for the three carbon sources, while at 24 h every carbon source showed a different pattern.

In sum, although the implication of *wspF* and *wspR* in biofilm formation through the secondary messenger c-di-GMP was evident, we argue that the differences in biofilm levels between all the mutants when we change the carbon source can be due to the particular regulation of other diguanylate cyclases present in the cytoplasm. Those enzymes could vary its activity or expression depending on cAMP balance, energy, oxidation levels and reduction power directly influenced by the carbon source (Huang *et al.*, 2013; Schirmer, 2016; da Costa Vasconcelos *et al.*, 2017). On the other hand, cyclase activity of WspR would likely differ from that of YedQ, as this protein comes from a different organism and they have different expression levels. This would also contribute to the differences in biofilm production between the *wspF* and *wspR* mutants and their transformation with the external cyclase and esterase.

### Colony morphology and swimming abilities

Colony morphology was observed in M9 agar plates with Congo Red and Coomassie staining for EPS inspection. Congo Red is a dye for extracellular polysaccharides, while Coomassie targets extracellular proteins. As a consequence, a colony with a deeper red colour usually correlates with higher biofilm forming ability. This assay revealed a very low red coloration for *P. putida wt* in all carbon sources tested, while instead revealing a coarse surface for *wspF* mutants in the citrate regime and the reddest colour in fructose as sole carbon source compared to the *wt* strain (Figure 3a). Although Congo Red and Coomassie staining is a qualitative assay for morphology description, and results do not always completely correlate to quantitative biofilm assays, observations for the *wspF* mutant were consistent with those shown in Figure 3. Swimming assays showed minimal halo width differences among mutants except in the case of wspF (Figure 3b), which was less able to move along the soft agar in all the tested carbon sources, forming smaller circles, a finding coherent with the higher amount of c-di-GMP produced by this strain.

**Figure 3:**
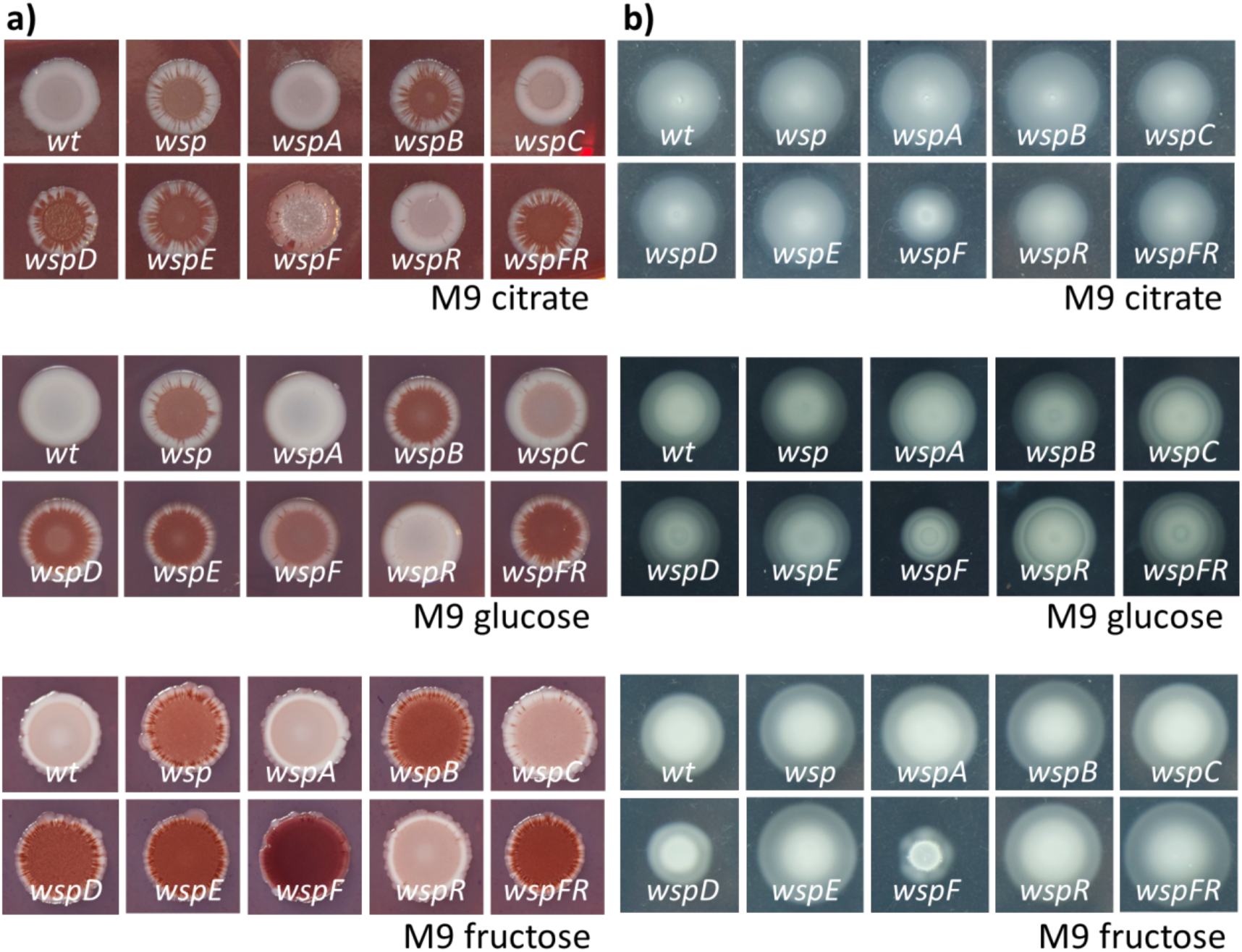
Colony morphology and swimming assay for *P. putida wsp* mutants. **(a)** EPS inspection of *wsp* mutants was carried out using Congo Red/Coomassie staining for M9 solid media supplemented with citrate, glucose or fructose. Colony colour intensity correlates with the amount of produced biofilm. **(b)** Swimming assays were performed in M9 0.3% agar (w/v) media supplemented with citrate, glucose or fructose as sole carbon sources. The *wspF* mutant was less able to swim in all of the carbon sources tested.

Taken together, these observations indicate that the *P. putida* Wsp complex works in a similar way as that of *P. aeruginosa*, i.e. the enzymatic activity of WspR regulates c-di-GMP levels of the cell under the control of WspF, and therefore also regulates biofilm levels. The influence of the carbon source in biofilm production is also remarkable. Some hypotheses and explanations to those considerations are addressed below.

### P. putida wsp complementation with P. aeruginosa’s genes

In order to see whether *Pseudomonas aeruginosa* Wsp proteins were able to complement *P. putida* mutants, we cloned the *wsp* operon, *wspF, wspR* or *wspF wspR* from *P. aeruginosa* PAO1 strain in a pSEVA238 plasmid (Silva-Rocha *et al.*, 2013) under the control of the XylS-*Pm* expression system. Then, we transformed the deletion mutants of *Pseudomonas putida* with the new plasmids harbouring *P. aeruginosa*’s pertinent gene and checked its phenotype both under induction of the expression system with 2.0 mM of 3MBz and at the basal level (Figure 4). Our main observation was that *wspF* from *P. aeruginosa* could complement the deficient *wspF P. putida* strain in terms of biofilm formation: the phenotype of the *wspF*^*PA*^ strain showed lower levels of biofilm than the non-complemented *wspF*, both when the expression of *wspF*^*PA*^ was induced and not induced (except in fructose at 6 hours when induced, which showed lower levels of biofilm than *wild type*). This result indicates that low levels of WspF from *P. aeruginosa* are enough to regulate WspR function from *P. putida*, as the XylS-*Pm* expression system has very low leakiness (Calles, B., in preparation). However, the rest of the complementations showed a higher biofilm activity than the *P. putida wt* strain, specially at 24 h and even when they were not induced under most of the conditions tested. These results let us infer that higher levels of WspR correlate with an increase in the amount of c-di-GMP and therefore biofilm formation as well. The possibility that *P. aeruginosa* WspR is more active than the corresponding enzyme in *P. putida* deserves further studies.

**Figure 4:**
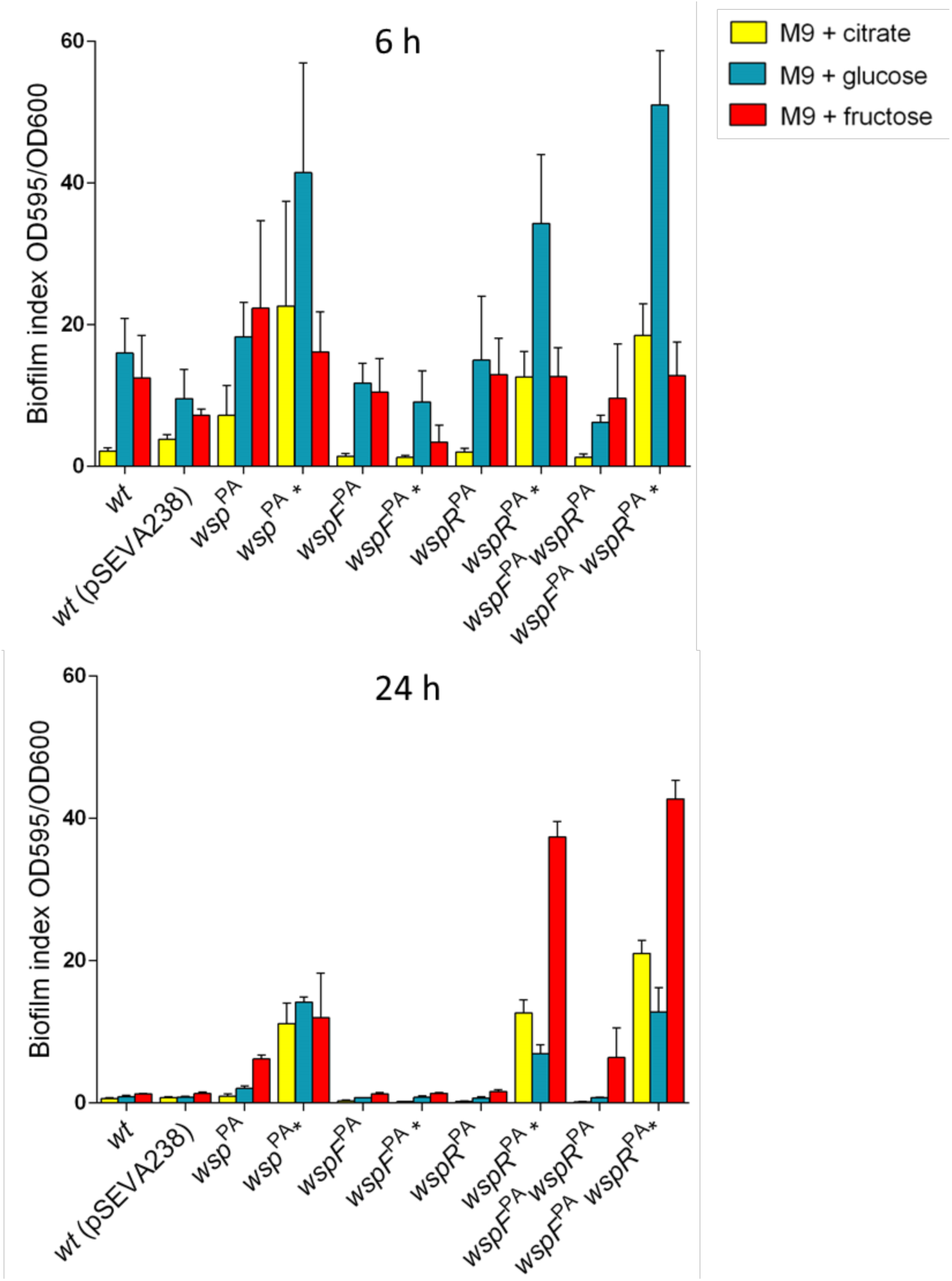
Biofilm assay for *P. putida wsp* mutants and their complementations with the orthologous genes from *P. aeruginosa*. The experiment was performed in M9 supplemented with citrate (yellow), glucose (blue) or fructose (red) media. *P. putida*’s *wsp, wspF, wspR* and *wspF wspR* mutants were transformed with pSEVA238 harbouring genes *wsp* (*wsp*^PA^), *wspF* (*wspF*^PA^), *wspR* (*wspR*^PA^) and *wspF wspR* (*wspF*^PA^ *wspR*^PA^) from *P. aeruginosa* respectively. *P. putida wt* was also transformed with the empty vector pSEVA238. The expression of complemented genes was analysed under non-inducing conditions or in the presence of 2 mM 3MBz as inducer of the system (indicated with an asterisk), in order to check the activity of *P. aeruginosa*’s genes under basal and high expression levels.

In the same work package we also ran colony morphology analysis and swimming ability tests of the complemented strains described in the section above, both under 2.0 mM 3MBz induction and without it (Supplementary Figure S1). Results showed that complementation of the *wspF* mutant strain with *wspF*^*PA*^ restored the wild type phenotype in citrate, and exhibited no important differences between high-expressed and low-expressed WspF^PA^. But in the other carbon sources, we observed a slight decrease of swimming ability under glucose and fructose regimes and small differences in colony morphology, when induced and not induced, compared to the *wild type*. On the contrary, a higher expression of Wsp^PA^, WspF^PA^WspR^PA^ and WspR^PA^ in their pertinent mutant background presented a higher red staining of EPS and a clear reduction of swimming capacity with haloes reduced to the minimum value possible. Combining these findings with the biofilm assays from Figure 3, it is possible that *P. aeruginosa* Wsp complex keeps a higher level of phosphorylated WspR, letting the c-di-GMP concentration rise and endowing the species with a greater biofilm-forming capacity.

### c-di-GMP produced by WspR and lifestyle regulators

Although c-di-GMP implication in biofilm formation is clear, this secondary messenger is also involved in multiple other functions (Romling *et al.*, 2013). Therefore, this molecule has to orchestrate a considerable number of activities through mechanisms other than gross variations in its levels. As data from different studies indicate, c-di-GMP cytoplasmic amounts do not differ homogenously when the appropriate stimulus occurs. This means that general c-di-GMP concentration remains the same, while some local variations are produced. Those confined alterations produce the desired response using specific modulators that respond to heterogeneous variations (Dahlstrom and O’Toole, 2017; Dahlstrom *et al.*, 2018). To shed some light on the way that c-di-GMP formed from WspR interacts with certain transcription factors and regulators to form biofilm in *P. putida*, the same type of experiments were performed using combined deletions of three regulators that respond to c-di-GMP in this context: for one side, we focused on FleQ and FleN, transcription factors that are clearly involved in the regulation of motility and sesility (Baraquet *et al*., 2012; Jimenez-Fernandez *et al*., 2016; Blanco-Romero *et al*., 2018). On a different side, FlgZ, also known as YgcR *E. coli*, is a protein that harbours a c-di-GMP recognizing PilZ domain, which has been determined as a motility regulator through certain flagellar parts (Martinez-Granero *et al*., 2014). We checked the biofilm forming phenotype of *flgZ, fleQ, fleN* and *fleQ fleN* deletions in addition to *wspF flgZ, wspF fleQ, wspF fleN* and *wspF fleQ fleN* associated deletions (Figure 5). On citrate supplemented media, we found after 6 and 24 h of incubation that each mutant for *fleQ, fleN* and *fleQ fleN* was very similar to its double mutant *fleQ wspF, fleN wspF* and *fleQ fleN wspF* respectively. This meant that the phenotype of *fleQ* mutant was dominant over *wspF*, and c-di-GMP formed in this last mutant could not translate to a biofilm production in the absence of *fleQ* gene. However, double mutant *flgZ wspF* behaved as simple mutant *wspF*, suggesting that *wspF* phenotype is dominant over *flgZ*. This means that c-di-GMP formed by WspR could form similar amounts of biofilm in the presence or the absence of FlgZ.

**Figure 5:**
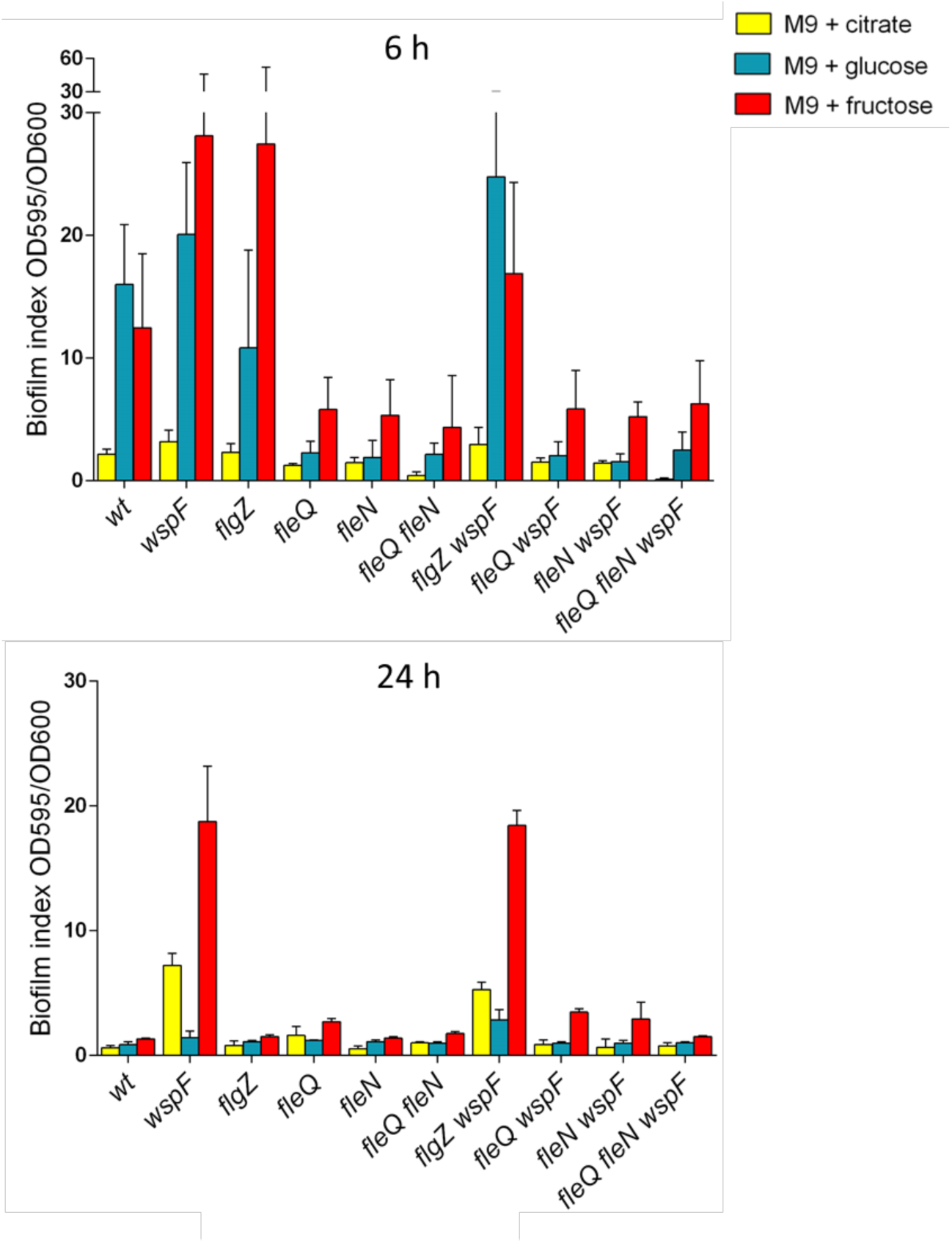
Biofilm assay of *P. putida* mutant FleQ, FleN and FlgZ in relation to WspF deficiency. Cultures of each of the mutants indicated were incubated in M9 medium supplemented with citrate (yellow), glucose (blue) and fructose (red) as sole carbon source during 6 h and 24 h. Simple deletions (*wspF, flgZ, fleQ* and *fleN*) were compared to double and triple mutants (*fleQ fleN, flgZ wspF, fleQ wspF, fleN wspF* and *fleQ fleN wspF*) in order to find epistatic relations.

When we checked the morphology phenotype of the various strains, we observed a common pattern in the *fleQ, fleN, fleQ fleN* mutants and same deletions in combination with *wspF* mutant (Supplementary Figure S2). The high levels of biofilm in the *wspF*-deleted strain dropped clearly when this mutation was combined with *fleQ, fleN* or *fleQ fleN* deletions, this last triple mutant being more evidently biofilm deficient at 6 h of incubation in citrate-supplemented media. However, only *fleQ* and *fleQ fleN* (but not *fleN*) mutants were impaired in swimming. Previous studies showed that FleQ is indispensable for this ability and for biofilm formation (Jimenez-Fernandez *et al.*, 2016), such that a *fleQ*-deficient bacteria are unable to swim or form biofilm independently of the cellular c-di-GMP concentration. Yet, in Congo Red and Coomassie staining, double mutant *wspF fleQ* had a phenotype more similar to the *wspF* mutant than to the *fleQ* mutant, suggesting a FleQ-independent mechanism for EPS secretion. On the other hand, the *flgZ wspF* double mutant showed a comparable phenotype to the *wspF* simple mutant, indicating that c-di-GMP formed by WspR did not affect motility or colony morphology through FlgZ in the conditions tested.

### Carbon source as a lifestyle modulator

As can be deduced from the results above, the carbon regime was clearly an influencing factor for final biofilm production. Differences in this production were probably caused by the point of the central metabolism where the carbon skeleton was added. Some previous works have noted the relevance of the kind of carbon source in biofilm formation (Reeves *et al*., 1996; Ramos *et al*., 2001; Pemmaraju *et al.*, 2016). In the case of *P. putida*, glucose enters the metabolism and has to go through a special glycolysis that resembles a mix of the Entner-Doudoroff (ED) pathway and the Pentose Phosphate pathway (PP), as it cannot be metabolized through the Tricarboxilic acid cycle (TCA) by the classical Embden-Meyerhof-Parnas (EMP) pathway (Nikel *et al.*, 2015). This regime led to intermediate levels of biofilm formation. On the other side, the use of fructose as a sole carbon source produced the highest levels of biofilm. Both hexoses can be channeled through ED to the PP for catabolism, but also for gluconeogenesis, which could be related with the higher EPS production of those conditions. Another interesting case is citrate, which can be incorporated directly to the TCA cycle and showed the lowest levels of biofilm staining, suggesting that this molecule was mainly used for catabolism.

### Conclusion

Along this work we have seen Wsp complex in *P. putida* works in a similar way than it does in *P. aeruginosa*. It regulates c-di-GMP levels of the bacteria through two main components: WspR and WspF. WspR is the protein that harbours the GGDEF domain, which endow it with the capacity to cycle c-di-GMP. WspF is a crucial protein for the regulation of WspR activity. For other side, the Wsp complex from *P. aeruginosa* is so evolutionary close to *P. putida*’s that WspF^PA^ can modulate WspR^PP^ phosphorylation by WspE^PP^ and slow down its cyclase activity, complementing the function of WspF^PP^ under certain conditions. The rest of complementations with *P. aeruginosa*’s genes showed different biofilm and swimming activities to its respective mutants, and although the *wild type* phenotype could not be rescued the reason was probably related to expression dose of the Wsp proteins or to a higher cyclase activity of WspR^PA^ compared to WspR^PP^. Further enzymatic activity studies should be done in order to test this last hypothesis. Regarding c-di-GMP effector proteins, transcription factor FleQ is a key regulator for biofilm production and swimming abilities of *P. putida*, so its deletion phenotype will be always dominant over other deletions of c-di-GMP forming or degrading proteins. Therefore, *fleQ* mutant and double mutant strains are impaired in those activities independently of the c-di-GMP levels. Finally, the considerable influence of carbon source on biofilm formation was clearly evident: *P. putida* and most of the mutants showed higher biofilm production levels with fructose as the only carbon source, while exhibiting an intermediate biofilm production in glucose and the least biofilm amounts in our experiments in citrate cultures. We consider the information obtained by those experiments is of relevance for further understanding of biofilm formation in *P. putida* and its possible control with multiple applications.

## EXPERIMENTAL PROCEDURES

### Strains, culture conditions and plasmid construction

Bacterial strains used in this study are listed in Table S1. All *P. putida* mutants were derived from the reference *wild type* strain KT2440 (Nelson *et al*., 2002). For the construction of new plasmids, *E. coli* strains DH5α, CC118, JM109, and DH5α λ*pir* were used. The last strain is necessary for the replication of R6K origin plasmids (as pEMG) because it encodes the *π* replication protein. *Pseudomonas aeruginosa* genes were cloned by PCR amplification using PAO1 strain as template. For general methods, *Escherichia coli* and *Pseudomonas putida* strains were cultured in Luria Bertani (LB) medium, while for the specific biofilm, swimming and colony morphology assays, M9 minimal medium supplemented with citrate 0.2%,(w/v), glucose 0.2% (w/v) or fructose 0.2% (w/v) was used. A temperature of 37°C was set for *E. coli* and *P. aeruginosa* cultures and of 30°C for *P. putida*. Solid plates were made with the same media adding 1.5% (w/v) for general incubations, while a concentration of agar at 0.3% (w/v) was utilized for swimming assays. Antibiotics were added at the proper concentrations when it was necessary: kanamycin (Km, 50 µg mL^-1^), ampicillin (Ap, 150 µg mL^-1^ for *E. coli* and 500 µg mL^-1^ for *P. putida*) and tetracycline (Tc, 10 µg mL^-1^ for *E. coli* and 5 µg mL^-1^ for *P. putida*). When needed, cultures were induced with the chemical 3-methilbenzoate (3MBz), also known as *m*-toluate (Sigma Aldrich, St Louis, Missouri, USA) at 2 mM. Plasmids built and employed in this work are listed in Table S2. Classical protocols of digestion, ligation and transformation for *E. coli* were used for the most of the constructs (Maniatis, 1982). PCRs were performed using Q5 High Fidelity polymerase or Phusion High Fidelity polymerase (both from New England Biolabs, Ipswich, Massachusetts, USA), and analytic PCRs were performed using MasterMix (Promega, Madison, Wisconsin, USA) or Green MasterMix (BioTools, Madrid, Spain). For digestions, restriction enzymes (New England Biolabs, Ipswich, Massachusetts, USA) were used and for ligations, T4 ligase (Roche, Basilea, Switzerland) was ordered. Plasmid DNA extraction from bacteria was performed using QIAprep Spin Miniprep Kit (Qiagen, Venlo, Netherlands). DNA was isolated from agarose bands with the NucleoSpin Extract II kit (MN, Düren, Germany). *E. coli* strains were either heat-shocked (90 seconds at 42 °C) or electro-shocked (cuvettes 0.2 cm, 2.4 kv) for transformations. For the transformations and deletions in *P. putida*, protocols for Gram-negative bacteria described by Martinez-Garcia and de Lorenzo (2011) were used. For the clean deletions, pEMG plasmids (listed in Supplementary Table S2) were built with the 500 bp’s flanking regions upstream and downstream of the genes to be deleted. Primers used for the amplification of this homology regions can be found in Supplementary Table S3.

### Crystal Violet assays

The protocol described by O’Toole and Kolter (1998) was used to perform Crystal Violet experiments. Different *P. putida* mutant strains were cultured overnight (O/N) at 30°C in the medium of interest. The following morning, transparent 96-well plates with flat bottoms were inoculated using 0.2 µL of O/N preinocula in 200 µL of the same medium. The plates were placed at 30°C without shaking during 6 or 24 h, and then OD_600_ was measured again and the biofilm stained with Crystal Violet at 1% (w/v) and later dissolved with acetic acid 33% (w/v). Crystal Violet intensity was measured at OD_595_. Both OD_600_ and OD_595_ were measured using SpectraMax iD3 (Molecular Devices, California, USA) plate reader. Biofilm index was calculated by dividing both measurements OD_595_/OD_600_ and the resulting value was plotted.

### Swimming assays and Congo Red Coomassie tests

For the swimming experiments, M9 medium supplemented with the different carbon sources and 0.3% (w/v) agar was used. Overnight cultures grown in M9 supplemented with the corresponding carbon source were diluted to an OD_600_ of 0.5. Aliquots of 3 µL of every dilution were placed onto soft agar plates with the same medium. Pictures of the swimming halos were taken after 24 h of incubation at 30°C. Colony morphology of the different strains was studied adding 40 µg/µL of Congo Red and 20 µg/µL of Coomassie brilliant Blue on different M9 media. Overnight cultures were adjusted to OD_600_ of 0.5, and a 3 µL drop was placed on the top of the agar. The plates were incubated at 30°C during 7 days, following which pictures of the colonies were taken.

## ACKNOWLEDGEMENTS

This work was funded by the SETH Project of the Spanish Ministry of Science RTI 2018-095584-B-C42, MADONNA (H2020-FET-OPEN-RIA-2017-1-766975), BioRoboost (H2020-NMBP-BIO-CSA-2018), and SYNBIO4FLAV (H2020-NMBP/0500) Contracts of the European Union and the S2017/BMD-3691 InGEMICS-CM funded by the Comunidad de Madrid (European Structural and Investment Funds). Authors declare no conflict of interests.

## SUPPLEMENTARY MATERIALS

**Supplementary Table S1:** *Escherichia coli* and *Pseudomonas* strains used in this work

**Supplementary Table S2:** Plasmids built and used in this work

**Supplementary Table S3:** Primers designed for *wsp* experiments

**Supplementary Figure S1:** Colony morphology and swimming ability for *P. putida wsp* mutants and their complementation with orthologous genes from *P. aeruginosa*

**Supplementary Figure S2:** Colony morphology and swimming ability of strains lacking transcription factors FleQ and FleN and for the regulator FlgZ, associated to *wspF* mutant

