## Supplementary Materials for "The Wsp intermembrane complex mediates metabolic control of the swim-attach decision of *Pseudomonas putida*"

**Supplementary Table S1:** *Escherichia coli* and *Pseudomonas* strains used in this work.

| Strain | Description | Reference |
|---|---|---|
| **DH5α** | F$^-$, *supE44*, Δ*lacU169*, (φ80 *lacZDM15*), *hsdR17*, (*rkmk*$^+$), *recA1*, *endA1*, *thi1*, *gyrA*, *relA* | (Hanahan and Meselson, 1983) |
| **cc118** | F-, Δ(*ara-leu*) 7697, *araD*139, Δ (*lac*)X74, *phoA* Δ20, *ale*, *galK*, *thi*, *rpsE*, *rpoB* | (Manoil and Beckwith, 1985) |
| **JM109** | F′, *traD36*, *proA+B+*, *lacIq*, “(*lacZ*)M15/ “(*lac-proAB*), *glnV44* e14-, *gyrA96*, *recA1*, *relA1*, *endA1*, *thi*, *hsdR17* | (Yanisch-Perron *et al*., 1985) |
| **DH5α λpir** | λpir phage lysogen of DH5α, $\pi^+$ | Lab collection |
| **KT2440** | Prototrophic, wild-type strain derived of *P. putida* mt-2 without pWWO plasmid | (Nelson *et al.*, 2002) |
| **KT2440 Δ*wsp*** | KT2440 derivative with a full deletion of *wsp* operon | This work |
| **KT2440 Δ*wspA*** | KT2440 derivative with a full deletion of *wspA* | This work |
| **KT2440 Δ*wspB*** | KT2440 derivative with a full deletion of *wspB* | This work |
| **KT2440 Δ*wspC*** | KT2440 derivative with a full deletion of *wspC* | This work |
| **KT2440 Δ*wspD*** | KT2440 derivative with a full deletion of *wspD* | This work |
| **KT2440 Δ*wspE*** | KT2440 derivative with a full deletion of *wspE* | This work |
| **KT2440 Δ*wspF*** | KT2440 derivative with a full deletion of *wspF* | This work |
| **KT2440 Δ*wspR*** | KT2440 derivative with a full deletion of *wspR* | This work |
| **KT2440 Δ*wspFR*** | KT2440 derivative with a full deletion of *wspFR* | This work |
| **KT2440 Δ*flgZ*** | KT2440 derivative with a full deletion of *flgZ* | This work |
| **KT2440 Δ*fleQ*** | KT2440 derivative with a full deletion of *fleQ* | This work |
| **KT2440 Δ*fleN*** | KT2440 derivative with a full deletion of *fleN* | This work |
| **KT2440 Δ*fleQfleN*** | KT2440 derivative with full deletions of *fleQ* and *fleN* | This work |
| **KT2440 Δ*wspFflgZ*** | KT2440 derivative with full deletions of *wspF* and *flgZ* | This work |
| **KT2440 Δ*wspFfleQ*** | KT2440 derivative with full deletions of *wspF* and *fleQ* | This work |
| **KT2440 Δ*wspFfleN*** | KT2440 derivative with full deletions of *wspF* and *fleN* | This work |

| | | |
|---|---|---|
| **KT2440 Δ*wspFfleQfleN*** | KT2440 derivative with full deletions of *wspF, fleQ* and *fleN* | This work |
| **PAO1** | *Pseudomonas aeruginosa* wild-type strain | (Stover *et al.*, 2000) |

**Supplementary Table S2:** Plasmids built and used in this work.

| Plasmid | Description | Reference |
|---|---|---|
| **pEMG** | Km$^R$, *ori*R6K, suicide plasmid with two I-SceI sites flanking the *lacZα* polylinker | (Martinez-Garcia and de Lorenzo, 2011) |
| **pEMGwsp** | Km$^R$, *ori*R6K, two 500 bp flanking fragments of *wsp* operon from *P. putida* | This work |
| **pEMGwspA** | Km$^R$, *ori*R6K, two 500 bp flanking fragments of *wspA* gene from *P. putida* | This work |
| **pEMGwspB** | Km$^R$, *ori*R6K, two 500 bp flanking fragments of *wspB* gene from *P. putida* | This work |
| **pEMGwspC** | Km$^R$, *ori*R6K, two 500 bp flanking fragments of *wspC* gene from *P. putida* | This work |
| **pEMGwspD** | Km$^R$, *ori*R6K, two 500 bp flanking fragments of *wspD* gene from *P. putida* | This work |
| **pEMGwspE** | Km$^R$, *ori*R6K, two 500 bp flanking fragments of *wspE* gene from *P. putida* | This work |
| **pEMGwspF** | Km$^R$, *ori*R6K, two 500 bp flanking fragments of *wspF* gene from *P. putida* | This work |
| **pEMGwspR** | Km$^R$, *ori*R6K, two 500 bp flanking fragments of *wspR* gene from *P. putida* | This work |
| **pEMGwspFR** | Km$^R$, *ori*R6K, two 500 bp flanking fragments of *wspF*-*wspR* genes from *P. putida* | This work |
| **pEMGflgZ** | Km$^R$, *ori*R6K, two 500 bp flanking fragments of *flgZ* gene from *P. putida* | This work |
| **pEMGfleQ** | Km$^R$, *ori*R6K, two 500 bp flanking fragments of *fleQ* gene from *P. putida* | This work |
| **pEMGfleN** | Km$^R$, *ori*R6K, two 500 bp flanking fragments of *fleN* gene from *P. putida* | This work |
| **pSW-1** | Ap$^R$, *ori*RK2, *xylS*, bearing a *Pm→I-sce*I transcriptional fusion | (Wong and Mekalanos, 2000) |
| **pRK404A** | Tc$^R$, *ori*RK2, pRK290 derivative | (Ditta *et al.*, 1985) |
| **pYedQ** | Tc$^R$, *ori*RK2, pRK404A derivative, $p_{lac}$ →*yedQ* (formerly known as *yhcK*) diguanylate cyclase from *E. coli* | (Ausmees *et al.*, 2001) |
| **pYljH** | Tc$^R$, *ori*pBBR1, pBBR1-MCS3 derivative $p_{lac}$ | (Gjermansen *et al.*, |

| | | |
|---|---|---|
| | →*yljH* diguanylate esterase from *E. coli* | 2006) |
| **pSEVA238** | Km$^{R}$, *ori*pBBR1, *XylS-Pm* promoter, expression vector | (Silva-Rocha *et al.*, 2013) |
| **pWPA-W** | Km$^{R}$, *ori*pBBR1, *XylS-Pm* promoter, pSEVA238 where *wsp* operon from *P. aeruginosa* PAO1 was cloned with *XbaI/HindIII* | This work |
| **pWPA-F** | Km$^{R}$, *ori*pBBR1, *XylS-Pm* promoter, pSEVA238 where *wspF* from *P. aeruginosa* PAO1 was cloned with *XbaI/HindIII* | This work |
| **pWPA-R** | Km$^{R}$, *ori*pBBR1, *XylS-Pm* promoter, pSEVA238 where *wspR* from *P. aeruginosa* PAO1 was cloned with *XbaI/HindIII* | This work |
| **pWPA-FR** | Km$^{R}$, *ori*pBBR1, *XylS-Pm* promoter, pSEVA238 where *wspF* and *wspR* from *P. aeruginosa* PAO1 was cloned with *XbaI/HindIII* | This work |

**Supplementary Table S3:** Primers designed for *wsp* experiments. Restriction enzyme targets are marked in red and RBS's and their spacers are marked in blue.

| **Primer name** | **Sequence (5'→3')** | **TM** | **Constructed strain** |
|---|---|---|---|
| **wsp TS1F SacI** | TAGCTAGCAGAGCTCAGCAGCACAGCCAGGCGG | 62ºC | Δwsp/ΔwspA |
| **wsp TS1R** | CTCCAGGAGCaCAGGCGATGCAGCCCTCAGGAATTCACAAC | 62ºC | Δwsp |
| **wsp TS2F** | CATCGCCTGTGCTCCTGGAG | 62ºC | Δwsp/ΔwspR/ΔwspFR |
| **wsp TS2R XbaI** | TGCTACGCTCTAGAGGATGTCCAGGTAGGCGTTG | 62ºC | Δwsp/ΔwspR/ΔwspFR |
| **wspA TS1R** | CGCAGTTGCAAGTCGTTCATATTCAGCCCTCAGGAATTCAC | 58ºC | ΔwspA |
| **wspA TS2F** | ATGAACGACTTGCAACTGCG | 58ºC | ΔwspA |
| **wspA TS2R BamHI** | ATCGATCGATCGGATCCTTCATGTAGGCATCCCTGTC | 58ºC | ΔwspA |
| **wspB TS1F SacI** | ATCGATCGATCGAGCTCATCGTCAAGGTTGCCGACC | 58ºC | ΔwspB |
| **wspB TS1R** | GAAGAAACGCTGTTCGTTCATCGCGGTCAGACTTTGAAGC | 58ºC | ΔwspB |
| **wspB TS2F** | ATGAACGAACAGCGTTTCTTC | 58ºC | ΔwspB |
| **wspB TS2R BamHI** | ATCGATCGATCGGATCCCTTCGTCCAGAGCATCGAAG | 58ºC | ΔwspB |
| **wspC TS1F SacI** | ATCGATCGATCGAGCTCATGAACGACTTGCAACTGCG | 58ºC | ΔwspC |
| **wspC TS1R** | GCATCGCGAATTCACTGTTCTTCATGTAGGCATCCCTGTC | 58ºC | ΔwspC |
| **wspC TS2F** | GAACAGTGAATTCGCGATGC | 58ºC | ΔwspC |
| **wspC TS2R BamHI** | ATCGATCGATCGGATCCGTCGATTTCCTCGACAGCC | 58ºC | ΔwspC |
| **wspD TS1F SacI** | ATCGATCGATCGAGCTCTATGTGCGTCACGAAGGCG | 58ºC | ΔwspD |
| **wspD TS1R** | CGCATTTGCTCTGGGGTCATCGCGAATTCACTGTTCATCG | 58ºC | ΔwspD |
| **wspD TS2F** | ATGACCCCAGAGCAAATGCG | 58ºC | ΔwspD |

| | | | |
|---|---|---|---|
| **wspD TS2R BamHI** | ATCGATCGATCGGATCCGCGCTTGCCAGCCTTGCG | 58ºC | ΔwspD |
| **wspE TS1F SacI** | ATCGATCGATCGAGCTCCGTTTGCTCGACCGCTATG | 58ºC | ΔwspE |
| **wspE TS1R** | CCCTGAGCAACTCCGATCAGTCATGACAGGCTCCGCTGC | 58ºC | ΔwspE |
| **wspE TS2F** | CTGATCGGAGTTGCTCAGGG | 58ºC | ΔwspE |
| **wspE TS2R BamHI** | ATCGATCGATCGGATCCGCAAACCCTTGAGCAGCAC | 58ºC | ΔwspE |
| **wspF TS1F EcoRI** | GCCTATGGAATTCCCTTGCTCGATGATGGCTC | 57ºC | ΔwspF/ ΔwspFR |
| **wspF TS1R** | GGCGTGATTCAATACATTTCGTTCATCCCTGAGCAACTCCG | 57ºC | ΔwspF |
| **wspF TS2F** | CGAAATGTATTGAATCACGCC | 57ºC | ΔwspF |
| **wspF TS2R BamHI** | CGGTACTAGGATCCGCACGGTAGGCCTCGTCG | 57ºC | ΔwspF |
| **wspR TS1F SacI** | TAGCTAGCAGAGCTCGCCATCGTGCTGGTCCAGC | 62ºC | ΔwspR |
| **wspR TS1R** | CTCCAGGAGCaCAGGCGATGGGCGTGATTCAATACATTTCG | 62ºC | ΔwspR |
| **wspFR TS1R** | CTCCAGGAGCaCAGGCGATGTTCATCCCTGAGCAACTCCG | 60ºC | ΔwspFR |
| **wsp Del check F** | GCAAGTACGGTGGCAAGGC | 58ºC | ΔwspFR |
| **FleQ TS1F SacI** | ATCGATCGATCGAGCTCCCTCCGCCGAGCAATAATACC | 58ºC | ΔfleQ |
| **FleQ TS1R** | GAACACCCAGCCCAAACATCGCAATAGCAACTTCCCTAG | 58ºC | ΔfleQ |
| **FleQ TS2F** | GATGTTTGGGCTGGGTGTTC | 58ºC | ΔfleQ |
| **FleQ TS2R BamHI** | ATCGATCGATCGGATCCGACCATCGATCACGATCAC | 58ºC | ΔfleQ |
| **FleN TS1F SacI** | ATCGATCGATCGAGCTCATCGATACTGCCGGCCTGC | 58ºC | ΔfleN |
| **FleN TS1R** | CATCTTGAAGCCGCTGGCGTGTCTGTTCTTTACCTTGTCTC | 58ºC | ΔfleN |
| **FleN TS2F** | ACGCCAGCGGCTTCAAGATG | 58ºC | ΔfleN |
| **FleN TS2R** | ATCGATCGATCGGATCCCGCGCAGG | 58ºC | ΔfleN |

| | | | |
|---|---|---|---|
| **BamHI** | TTCGACCTGGC | | |
| **YcgR TS1F SacI** | TAGCTAGCAGAGCTCATGACATTGC CCCGATGCAGG | 58ºC | ΔycgR |
| **YcgR TS1R** | GCGTTTTCTGTTTTACCGACCGCTTA TTGTTCTCCAGGCAAAAGC | 58ºC | ΔycgR |
| **YcgR TS2F** | CGGTCGGTAAAACAGAAAACG | 58ºC | ΔycgR |
| **YcgR TS2R BamHI** | CGGTACTAGGATCCAGTTCTGCTTGT ACCCGCTGG | 58ºC | ΔycgR |
| **Wsp PA F XbaI** | CGGGGATCCTCTAGAAGGAGGAAAA ACATATGAAGAACTGGACTGTTCGC CAGC | 60ºC | pWPA-W |
| **Wsp PA R HindIII** | GTTTTCCCAGTCACGACGCGGCCGC AAGCTTGTACGCCGGCCTCTATTTAA TG | 60ºC | pWPA-W |
| **WspF PA F XbaI** | CGATCGATCTATCTAGAAGGAGGAA AAACATATGAGGATCGGAATCGTCA ATG | 60ºC | pWPA-F/ pWPA-FR |
| **WspF PA R HindIII** | TTCGATCTTAGCAAGCTTCTAATCGA ATACCTCCGCCA | 60ºC | pWPA-F |
| **WspR PA F XbaI** | CGATCGATCTATCTAGAAGGAGGAA AAACATATGCACAACCCTCATGAGA GC | 60ºC | PWPA-R |
| **WspR PA R HindIII** | TTCGATCTTAGCAAGCTTGTACGCCG GCCTCTATTTAATG | 60ºC | PWPA-R/ pWPA-FR |

**Supplementary Figure S1:** Colony morphology and swimming ability for *P. putida wsp* mutants and their complementation with orthologous genes from *P. aeruginosa*.

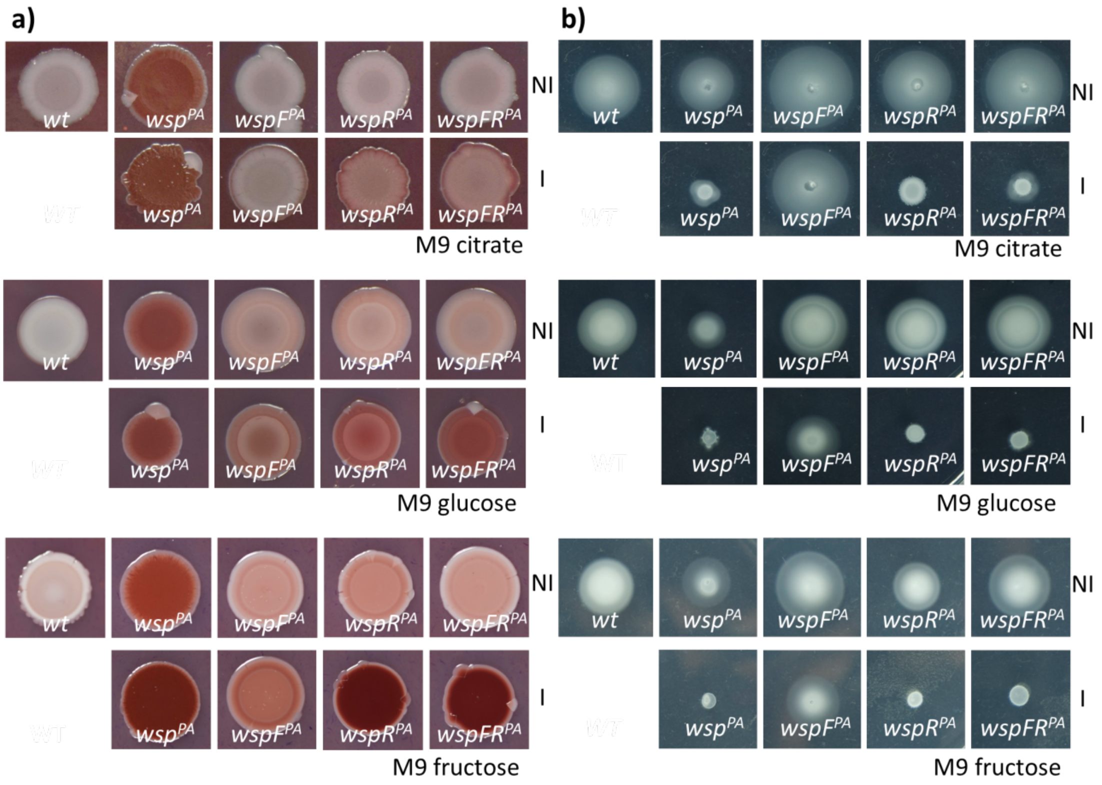


**(a)** Colony morphology and EPS accumulation was analysed by Congo Red and Coomassie staining in M9 agar supplemented with citrate, glucose or fructose as sole carbon sources. Inspection of complemented strains was done under non-induced (NI) and induced (I) conditions to test the influence of protein levels. **(b)** Swimming assay performed in M9 0.3% agar (w/v) supplemented with citrate, glucose and fructose. Again, Genes provided to complement the corresponding mutation were induced (I) and non-induced (NI) to check whether protein levels have an influence on phenotypes.

**Supplementary Figure S2:** Colony morphology and swimming ability of strains lacking transcription factors FleQ and FleN and for the regulator FlgZ, associated to *wspF* mutant.

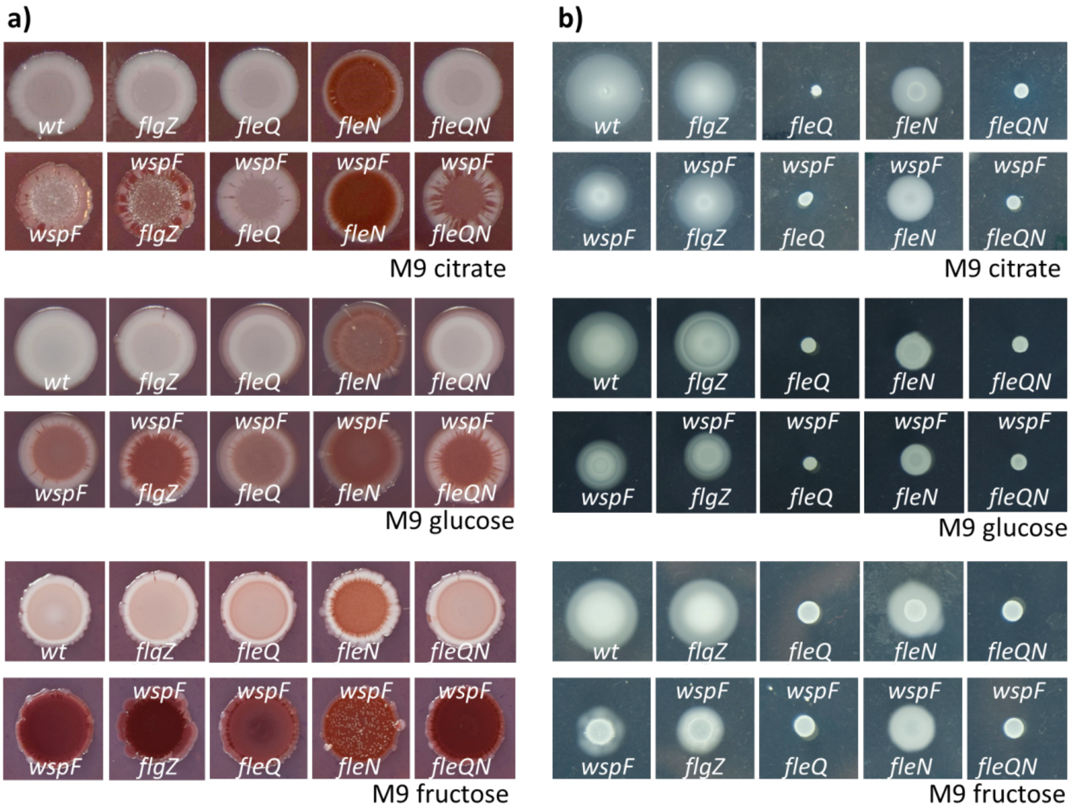


**(a)** Colony morphology and EPS levels were checked using Congo Red/Coomassie staining in M9 agar media supplemented with citrate, glucose or fructose. **(b)** Swimming assay for the same strains was performed in M9 0.3% agar (w/v) media supplemented with citrate, glucose or fructose as sole carbon sources for the different mutants.
